# MoCpa1-mediated arginine biosynthesis is crucial for fungal growth, conidiation, and plant infection of *Magnaporthe oryzae*

**DOI:** 10.1101/2020.12.27.424512

**Authors:** Osakina Aron, Min Wang, Anjago Wilfred Mabeche, Batool Wajjiha, Shuai Yang, Haixia You, Zonghua Wang, Wei Tang

## Abstract

Arginine is an important amino acid involved in processes such as cell signal transduction, protein synthesis, and sexual reproduction. To understand the biological roles of arginine biosynthesis in pathogenic fungi, we used Cpa1, the carbamoyl phosphate synthase arginine-specific small chain subunit in *Saccharomyces cerevisiae* as a query to identify its ortholog in *Magnaporthe oryzae* genome database and named it MoCpa1. MoCpa1 is a 471-amino acid protein containing the CPSase_sm_chain domain and the GATase domain. MoCpa1 transcripts were highly expressed at the conidiation, early-infection, and late-infection stages of the fungus. Targeted deletion of *MoCPA1* gene resulted in the Δ*Mocpa1* mutant exhibiting arginine auxotrophy on MM, confirming its role in *de novo* arginine biosynthesis. The Δ*Mocpa1* mutant presented significantly decreased sporulation with some of its conidia being defective in morphology. Furthermore, the Δ*Mocpa1* mutant was nonpathogenic on rice and barley leaves, which was a result of defects in appressorium-mediated penetration and restricted invasive hyphal growth within host cells. Addition of exogenous arginine partially rescued conidiation and pathogenicity defects on the barley and rice leaves, while introduction of *MoCPA1* gene in Δ*Mocpa1* mutant fully complemented the lost phenotype. Further confocal microscopy examination revealed that MoCpa1 is localized in the mitochondria. In summary, our results demonstrate that MoCpa1-mediated arginine biosynthesis is crucial for fungal development, conidiation, appressorium formation and infection-related morphogenesis in *M. oryzae*, thus serving as an attractive target for mitigating obstinate fungal plant pathogens.

## INTRODUCTION

Rice (*Oryza sativa*) is one of the major sources of food consumed by humans; however, rice yields continue to be under threat from the rice blast disease caused by *Magnaporthe oryzae*, which poses a great challenge to global food security. Due to its economic significance and genetic tractability, the blast fungus has been developed as a model organism for plant-fungus interaction studies (Talbot, 2003). Disease onset occurs when an asexual spore germinates and develops a specialized dome-shaped structure called an appressorium upon landing on the rice plant surface (Dagdas et al., 2012). The mature appressorium then accumulates enormous turgor pressure (8 MPa) that helps it puncture the rice cuticle, thus facilitating its entry into plant cells (Howard, Ferrari, Roach, & Money, 1991). While inside the host cell, the fungus differentiates into bulbous invasive hyphae (IH), which spread to the adjacent plant cells. The infection dynamics of *M. oryzae* involve an initial hemibiotrophic strategy that lasts approximately 4-6 days, during which it colonizes the living host cells without causing damage to the host before entering into a devastating necrotrophic phase where the fungus rapidly destroys the infected host tissue (Kankanala, Czymmek, & Valent, 2007). Later, the aerial conidiophore differentiates from the hyphae of the infected leaves and releases new conidia that act as inocula for secondary infection.

Amino acids are important biomolecules that contribute to various functions ranging from protein synthesis, cell growth, development, and energy production. Numerous studies have demonstrated that amino acid-related metabolic processes are crucial for growth and virulence in pathogenic fungi (Seong, Hou, Tracy, Kistler, & Xu, 2005; Takahara, Huser, & O’Connell, 2012). In *M. oryzae*, acetolactate synthase subunits (ALS) MoIlv2 and MoIlv6 were found to catalyze the first step in leucine, isoleucine, and valine biosynthesis, whereas threonine deaminase (MoIlv1) was found to be involved in isoleucine biosynthesis. Deletion of MoIlv2, Mollv6, and MoIlv1 resulted in defects in conidiation, appressorial penetration, and pathogenicity (Du et al., 2014; Du et al., 2013). Cystathionine beta-lyase (MoStr3) and methylene tetrahydrofolate reductase (MoMet13) were shown to be critical for methionine biosynthesis, and their respective deletion mutants exhibited defects in aerial hyphal growth, melanin pigmentation, and pathogenicity (Wilson et al., 2012; Yan et al., 2013). An aminoadipate reductase (MoLys2) was shown to be crucial for the development and pathogenicity of *M. oryzae* by modulating lysine biosynthesis (Y. Chen et al., 2014). Recently, isopropylmalate isomerase (MoLeu1), which orchestrates leucine biosynthesis, was determined to be important in fungal development and pathogenicity in *M*. *oryzae* (Tang et al., 2019).

Arginine is a semi-essential amino acid that is involved in an array of processes, such as cell, signal transduction, protein synthesis, and sexual reproduction (Bedford & Richard, 2005; Wu et al., 2009). Moreover, it has been shown that arginine, through nitric oxide synthetase (NOS) activity, is a precursor of nitric oxide (NO) in animals (Appleton, 2002). In fungi, NO generated from arginine has been reported to be crucial in conidium germination and appressorium formation (Prats, Carver, & Mur, 2008; J. Wang & Higgins, 2005).

The biosynthesis of arginine from glutamate involves a series of eight reaction steps, with the initial first five steps liberating ornithine. The ornithine generated reacts with carbamoyl phosphate via the ornithine carbamoyl transferase-catalyzed reaction, generating citrulline. Citrulline is further converted to argininosuccinic acid and ultimately to arginine by argininosuccinate lyase (Crabeel, Abadjieva, Hilven, Desimpelaere, & Soetens, 1997; Cunin, Glansdorff, Pierard, & Stalon, 1986; Slocum, 2005). Although previous studies have shown that arginine biosynthesis is important for fungal development and pathogenicity in *M. oryzae* (Xinyu Liu, Cai, et al., 2016; Y. Zhang et al., 2015), direct evidence demonstrating the involvement of MoCpa1 in remains unclear. Here, we identified and characterized the role of MoCpa1 in rice blast fungus, we established that deletion *MoCPA1* impaired with arginine biosynthesis and compromised fungal developmental processes such as aerial hyphae formation, conidiation, appressorium formation and penetration and pathogenicity. This study provides additional evidence on the importance of amino acid biosynthetic processes in the development and pathogenicity of fungal pathogens, and serves as a suitable target for combating obstinate fungal pathogens

## RESULTS

### Identification of MoCpa1 in *M. oryzae*

To identify *M. oryzae* carbamoyl phosphate synthetase small subunit, the amino acid sequence of Cpa1 from *Saccharomyces cerevisiae* was used for a blastP search in Kyoto Encyclopedia of Genes and Genome (KEGG) section for *M. oryzae* (http://www.kegg.jp/kegg-bin/show_organism?org=mgr). The obtained MoCpa1 protein sequence (MGG_01743), consisted of 471 amino acid and it was used for Pfam-based domain prediction analysis. Results obtained showed that MoCpa1 contained CPSase_sm_chain and GATase domains (Fig. 1A). To established whether these domains are present in other fungi including *Saccharomyces cerevisiae* (Sc), *Sclerotinia sclerotiorum* (Ss), *Fusarium oxysporum* (Fo), *Neurospora crassa* (Nc), *Fusarium graminearum* (Fg), *Trichoderma reesei* (Tr), *Ustilago maydi*s (Um). Their respective Cpa1 protein sequence were retrieved from National Center for Biotechnology Information (NCBI) https://www.ncbi.nlm.nih.gov and were used to perform additional Pfam-based domain prediction. Results obtained revealed that the CPSase_sm_chain and GATase domains are conserved among the Cpa1 orthologs in fungi (Fig. 1A). Phylogenetic analysis of the domains showed that the MoCpa1 CPSase_sm_chain domain is closely related to that of *Neurospora crassa* (Fig. 1B), while the MoCpa1 GATase is closer to that of *Trichoderma reesei* (Fig. 1C).

**Fig 1.**
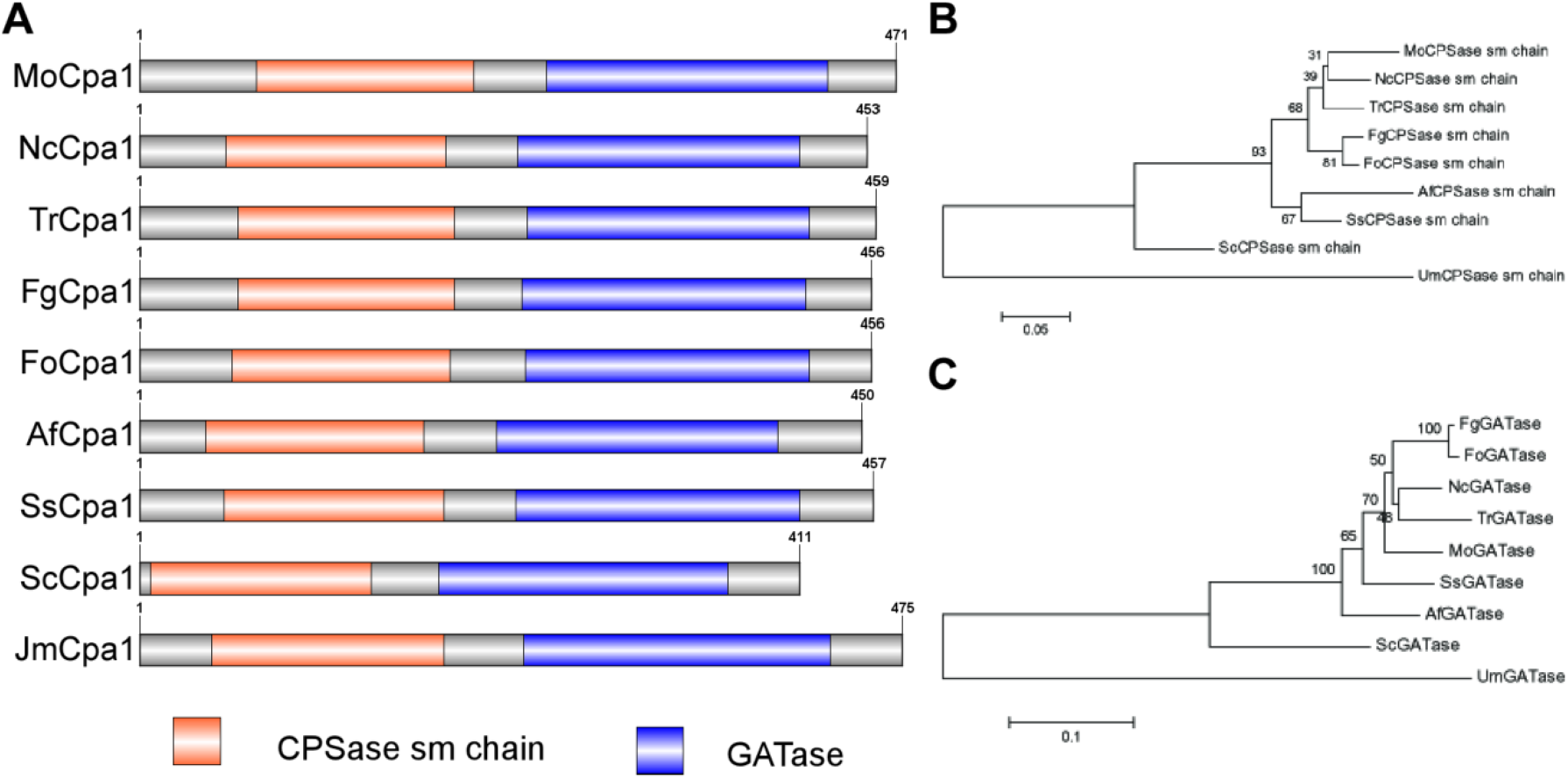
Domain architecture and phylogeny of MoCpa1 and Cpa1 orthologs in different fungal groups. (**A**) Represents the domain architecture showing the MoCpa1 CPSase_sm_chain and GATase domains and the Cpa1 CPSase_sm_chain and GATase amongst the other fungi. **(B**) Representation of the Maximum Likelihood phylogenetic outlook for the CPSase domain in fungi. (**C**) Phylogeny representing of the Maximum Likelihood outlook for GATase domain in different fungi. The Maximum Likelihood was tested with 1000 bootstrap replicates

### Expression of *MoCPA1* at different developmental stages of *M. oryzae*

To gain insight into the role of *MoCPA1* in *M. oryzae*, we first used qRT-PCR analysis to examine the expression patterns of *MoCPA1* at various fungal developmental stages. We established an elevated expression of the MoCpa1 transcripts at the asexual, early-infection and late-infection stages of fungal development, this was in comparison with the mycelial stage (Fig. 2). We noted a 2.4-fold increase at the conidial stage and 2.8-fold, 2.6-fold, 3.8-fold and 4.0-fold increases at 8, 24, 48, and 72 hours after inoculation respectively (Fig. 2). These results indicated that *MoCPA1* likely plays a key role in both asexual development and infection in *M. oryzae.* To determine the exact roles of *MoCPA1*, we generated Δ*Mocpa1* deletion mutant by replacing the entire *MoCPA1* coding region from Guy11 with the hygromycin resistance (HPH) gene. A total of two putative *MoCPA1* gene deletion transformants were obtained, which were subsequently confirmed by Southern blot analysis (Fig S1).

**Fig. 2.**
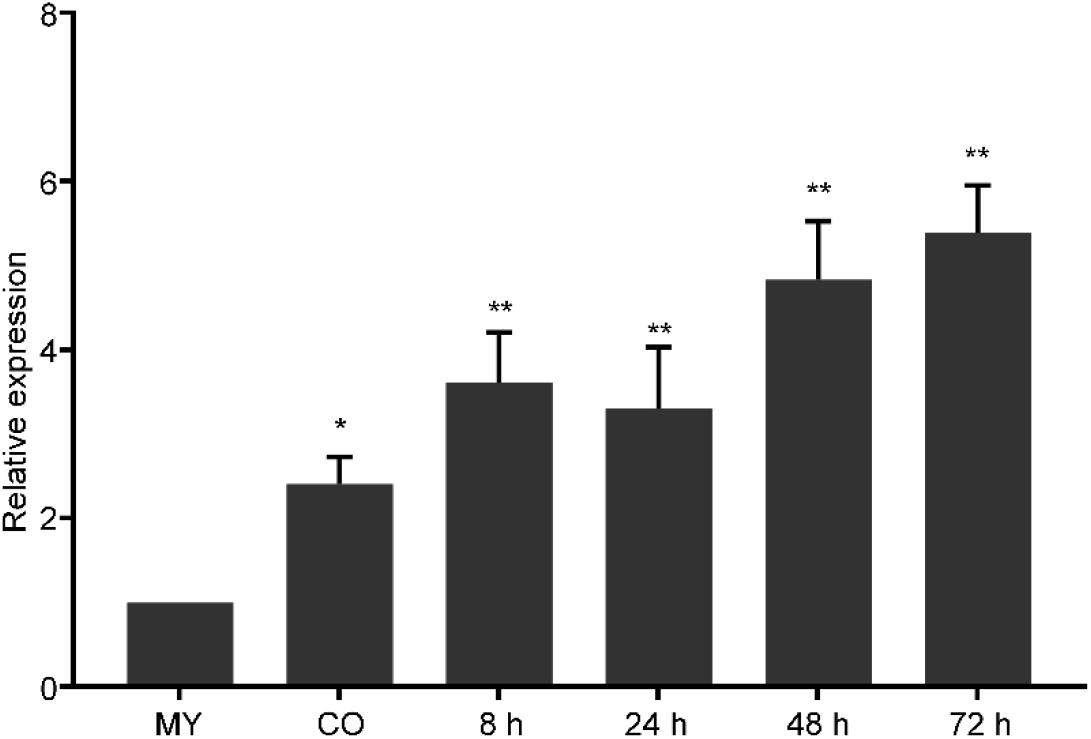
Expression patterns of *MoCPA1* at different developmental stages. RNA was extracted from mycelia (MY), conidia (CO), and rice leaves inoculated with conidia from (5×10^4^ spores/mL) at 8, 24, 48, and 72 h, respectively. The *ACTIN* gene (MGG_03982) was used for normalization, the expression level of *MoCPA1* at the mycelial stage was considered 1 for further comparisons. Data was analyzed with GraphPad-prism6, the error bars represents the standard deviation and the asterisks (\**P*<0.05) and (\*\**P*<0.01) represent significance deference in expression based on one way-ANOVA

### *MoCPA1* is indispensable for development of aerial hyphae and arginine biosynthesis

To assess the contribution of MoCpa1 in vegetative growth of *M. oryzae*, we cultured the Δ*Mocpa1* mutant, WT, and the complemented strains on CM, SDC, MM, and OTM solid media for 8 days at 26°C and measured their colony diameter. After 8 days of incubation on CM medium, we established no significant difference in colony diameter and hyphae morphology of the Δ*Mocpa1*, WT, and the complemented strains. However, the Δ*Mocpa1* formed very thin colonies on SDC and OTM due to reduced aerial hyphae in contrast to the thick colonies formed by WT and the complemented strain (Fig. 3A). The impaired development of aerial hyphae observed with Δ*Mocpa1* prompted us to speculate a possibility of Δ*Mocpa1* mutant undergoing hyphal autolysis; therefore, we analyzed the expression of autolysis-related genes in Δ*Mocpa1* by qRT-PCR. The results obtained showed that the expression of these genes increased in the Δ*Mocpa1* mutant (Fig S2).Since Δ*Mocpa1* mutant showed attenuated growth on MM medium, we next sort to establish whether MoCpa1 could be involved in arginine biosynthesis. We cultured Δ*Mocpa1* on MM plates supplemented with various concentrations of arginine (0.1 mM, 0.625 mM, 1.25 mM, 2.5 mM, and 5 mM). Our data revealed that exogenous arginine rescued the growth defects of the Δ*Mocpa1* deletion mutant, with faster growth, and increased aerial hyphae formation recorded at a relatively high concentration of arginine (Fig. 3C). These findings implied that the deletion of *MoCPA1* interfered with arginine biosynthesis leading to arginine deficiency and consequently resulting in impaired aerial hyphal development in the Δ*Mocpa1* deletion mutant.

**Fig. 3.**
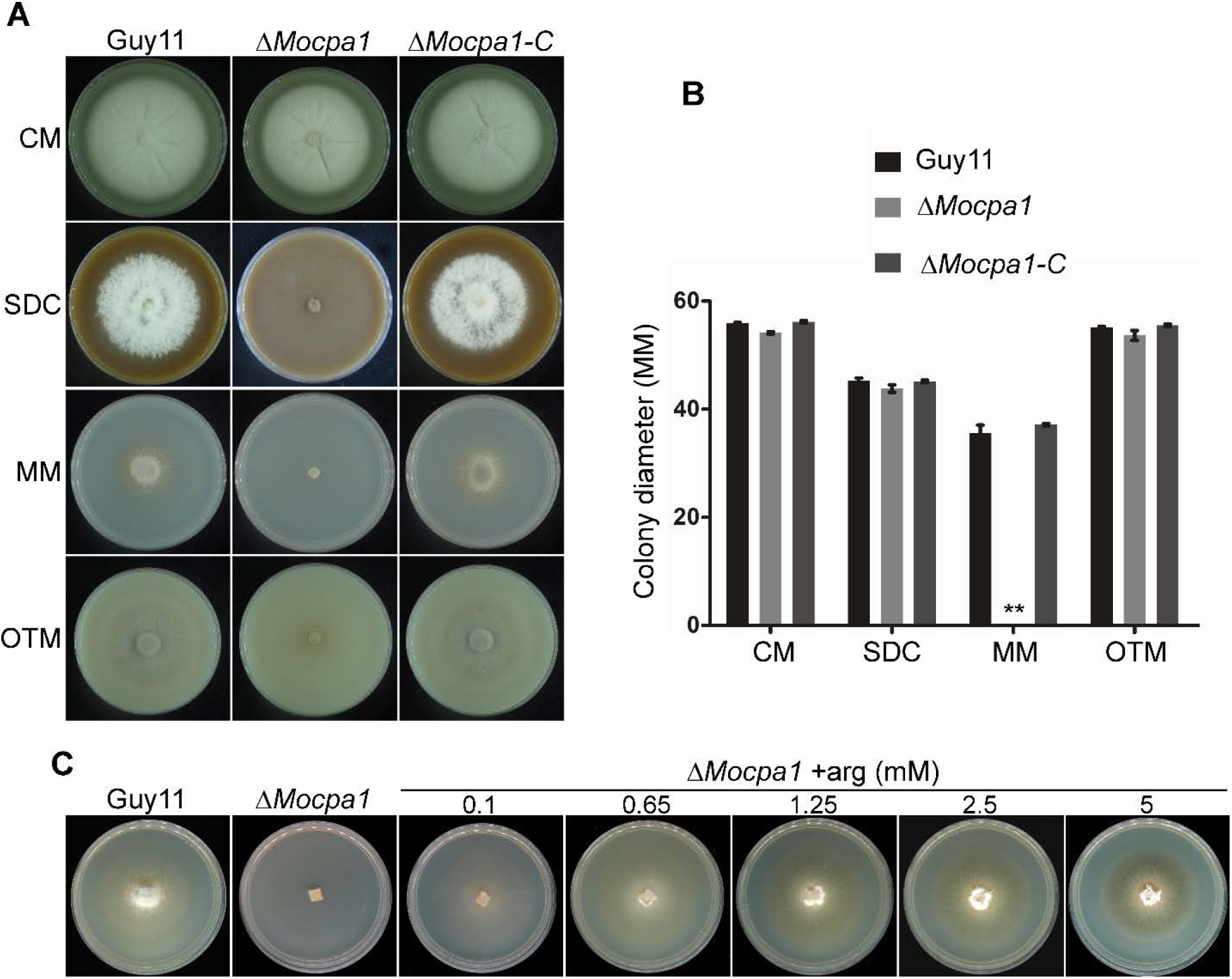
Δ*Mocpa1* exhibited a defect in hyphal growth. (**A**) Colony morphology and diameter of Δ*Mocpa1*, WT, and complemented strain cultured on CM, SDC, MM, and OTM for 8 days. (**b**) Statistical representations of average growth characteristics of Δ*Mocpa1* mutant compared to Guy11 and the complemented strain on CM, SDC, MM, and OTM for 8 days. Data used for statistical analysis were obtained from three independent biological experiments consisting of three replicates each time. One-way anova (nonparametric) statistical analysis was applied using graphpad prism6. Error bars represent the standard deviation while asterisk represent a significant difference in growth between Δ*Mocpa1* mutant compared to Guy11 and the complemented strain thus; asterisk (**p < 0.01). (**C**) Colony diameter of the Δ*Mocpa1* mutant cultured on MM supplemented with various concentrations (0.1 mm, 0.65 mm, 1.25 mm, 2.5 mm, and 5 mm) of arginine (arg) for 8 days at 28°c. Growth was compared to the Guy11 wild type strain on MM without exogenous arginine.

### MoCpa1 contributes to asexual reproduction in *M. oryzae*

In *M. oryzae*, asexual spores (conidia) play an important role in the disease cycle of rice blast (Wilson & Talbot, 2009). To determine the role of the *MoCPA1* gene in asexual sporulation, we analyzed the conidiation of the Δ*Mocpa1*, WT, and complemented strain in detail by incubating them on rice bran media for 10 days. We observed fewer conidiophorenhad formed for the Δ*Mocpa1* mutant compared to WT and the complemented strain (Fig. 4a). Consistent with the reduced conidiophore development, we found that the Δ*Mocpa1* deletion mutant produced fewer conidia, with a 75% reduction in conidiation (Fig. 4b). Next we performed quantitative real-time PCR (qRT-PCR) analysis to examine the expression levels of genes important for conidiation and conidial morphogenesis and found that the expression of *MoCOS1*, *MoCOM1*, *MoCON6*, *MoCON7*, *MoHOX6*, and *MoHOX7* were notably reduced in expression in Δ*Mocpa1* (Fig. 4C), indicating that *MoCPA1* may regulate Conidiogenesis in *M. oryzae* through regulation of the expression of conidiation-related genes. These results therefore indicates that MoCpa1 is important for conidiation in rice blast fungus *M. oryzae*.

**Fig. 4.**
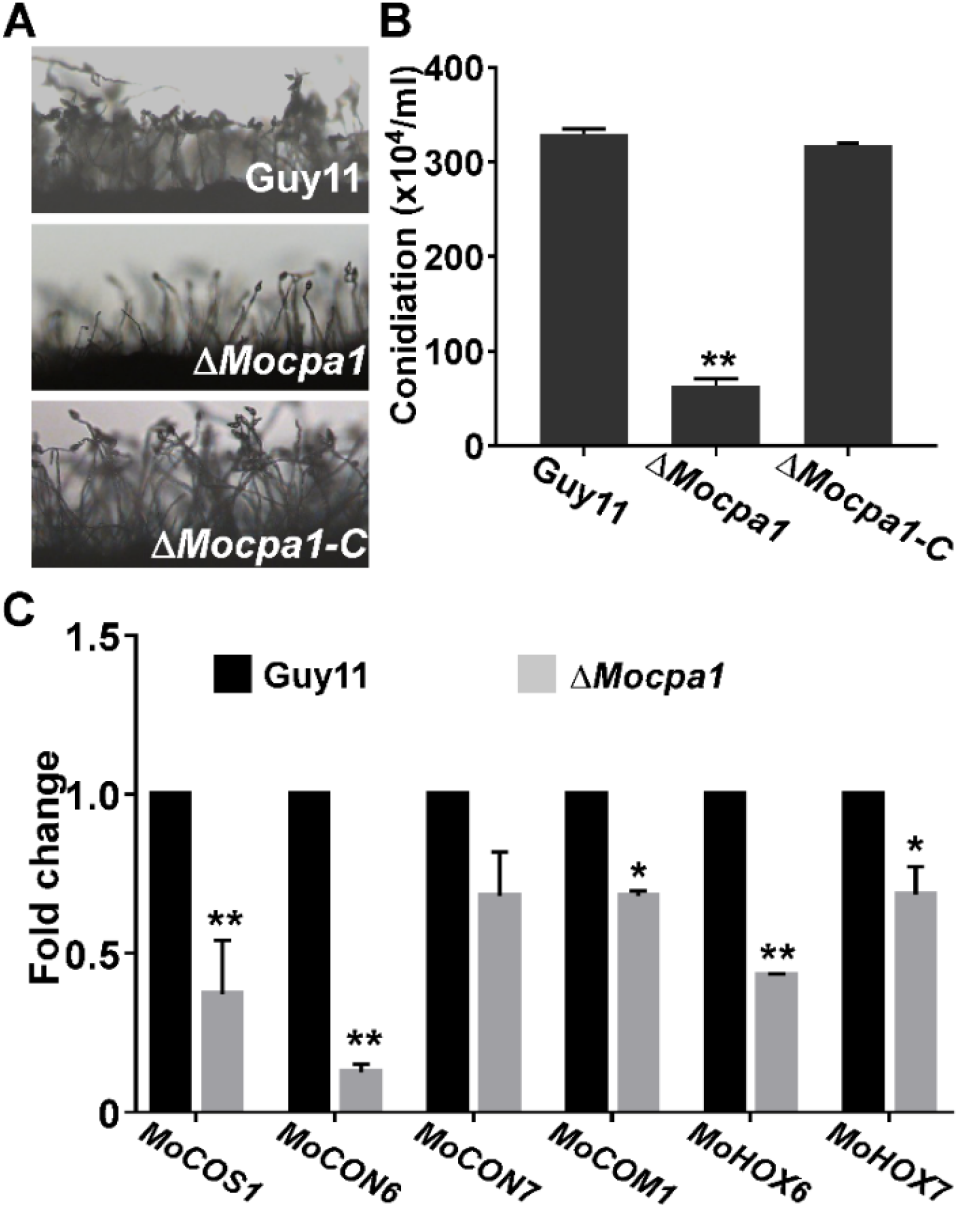
MoCpa1 contribute significantly to the conidiogenesis of *M. oryzae*. (A) Showed the distribution of conidiophore structures containing conidia produced by Δ*Mocpa1* mutant, the Guy11 wild type and the complementary strain (B) Statistical representation of the conidiation characteristics of Δ*Mocpa1* mutant, the Guy11 wild type and the complementary strain. Conidiation assays for Δ*Mocpa1* mutant, the Guy11 wild type and the complementary strain were evaluated from five independent biological repeats. The data was analyzed with GraphPad-prism6, error bars represent the standard deviation and the asterisks (**) represent significant differences (p < 0.01) according one – way ANOVA. **(B)** Expression patterns of conidiation related genes. RNA was extracted from mycelia of the WT and Δ*Mocpa1* mutant cultured in CM medium at 28°C after shaking for three days at 110 rpm. The ACTIN gene (MGG_03982) was used for normalization, the expression level of each gene tested in Guy11 at was considered 1 for comparisons. Data was analyzed with GraphPad-prism6, the error bars represents the standard deviation and the asterisks (*P<0.05) and (\*\**P*<0.01) represent significance deference in expression based on one way-ANOVA.

### MoCpa1 is crucial for conidial morphogenesis and appressorium formation

Rice blast fungus mainly produces normal three-cell conidia. Interestingly, we noted that in addition to the three-cell conidia, 10% of the conidia from the Δ*Mocpa1* mutant were 1-celled, while 40% constituted 2-celled (Fig. 5A and 5B). Since the appressoria form from conidia, we investigated whether all three different types of conidia produced by the Δ*Mocpa1* mutant could germinate and develop appressoria on an inductive surface. The results showed that at 4 h, 80% of Guy11 conidia produced appressoria, while only approximately 20% of Δ*Mocpa1* conidia formed appressoria. Eight hours later, 100% of Guy11 conidia had formed appressoria, while only approximately 40% of Δ*Mocpa1* conidia had formed appressoria (Fig. 5C and 5D). Further examination of appressorium formation at 12 h and 24 h did not yield any change in the number of appressoria formed by Δ*Mocpa1*, thus showing that deletion of *MoCPA1* compromised appressorium formation in the Δ*Mocpa1* mutant, with approximately 60% of its conidia unable to produce appressoria. The ability to form appressoria of the complemented strain was nearly the same as that of the WT. Taken together, these results indicate that *MoCPA1* is important for conidial morphogenesis and appressorium formation.

**Fig. 5.**
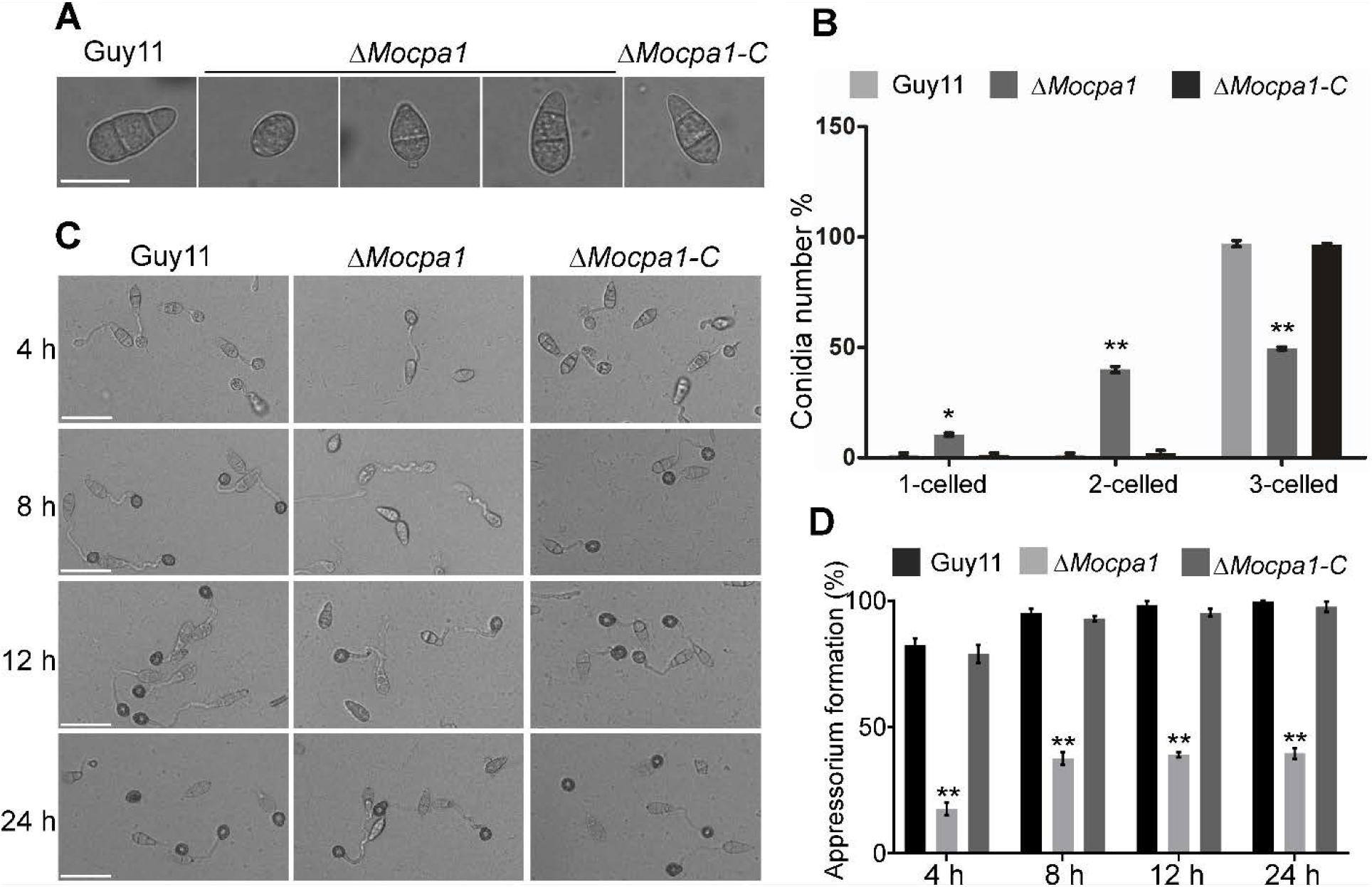
Defects in conidial morphogenesis and appressorium formation of the Δ*Mocpa1* mutant. (**A**) Conidia of Guy11, the Δ*Mocpa1* mutant, and the complemented strain were examined using differential interference contrast (DIC) microscopy. Bar=20 μm. (**B**) Statistics showing the percentage of conidium cell types for Guy11, the Δ*Mocpa1* mutant, and the complemented strain. Data analysis was by use of GraphPad-prism6, error bars represent the standard deviation, and the (p*< 0.05) and (p**< 0.01) represents significant differences according to ordinary one – way ANOVA (**C**) Appressoria of Guy11, the Δ*Mocpa1* mutant, and the complemented strain were examined at 4 h, 8 h, 12 h and 24 h on hydrophobic cover slips using differential interference contrast (DIC) microscopy. Bar=20 μm. (**D**) Statistics representing the percentage of appressorium formation on the hydrophobic cover slips for Guy11, the Δ*Mocpa1* mutant, and the complemented strain at 4 h, 8 h, 12 h and 24 h. The data was analyzed with GraphPad-prism6, error bars represent the standard deviation and the (p**< 0.01) represents significant differences according to ordinary one – way ANOVA.

### MoCpa1 is essential for full virulence in *M. oryzae*

To determine the contribution of *MoCPA1* in the virulence of *M. oryzae*, we conducted pathogenicity assay by spraying conidia from the Guy11, the Δ*Mocpa1* mutant, and the complemented strain onto 3-week-old rice seedlings. Seven days after inoculation, we noted that the rice seedlings sprayed with conidia from the wild-type Guy11 and the complemented strains showed pronounced lesions, while the rice seedlings challenged with conidia derived from Δ*Mocpa1* mutant did not show any disease symptoms (Fig. 6A). Moreover, we inoculated mycelial blocks and conidia from Guy11, the Δ*Mocpa1* mutant, and the complemented strain on both injured and intact barley leaves and monitored the disease progression. Similar results observed on rice leaves sprayed with strains conidia were replicated on both injured and intact barley leaves after 7 days of inoculation (Fig. 6B and 6C). These results therefore, shows that disruption of *MoCPA1* resulted in a complete loss of virulence of the Δ*Mocpa1* deletion mutant thus reinforcing the importance of MoCpa1 in the pathogenicity of *M. oryzae*.

**Fig. 6.**
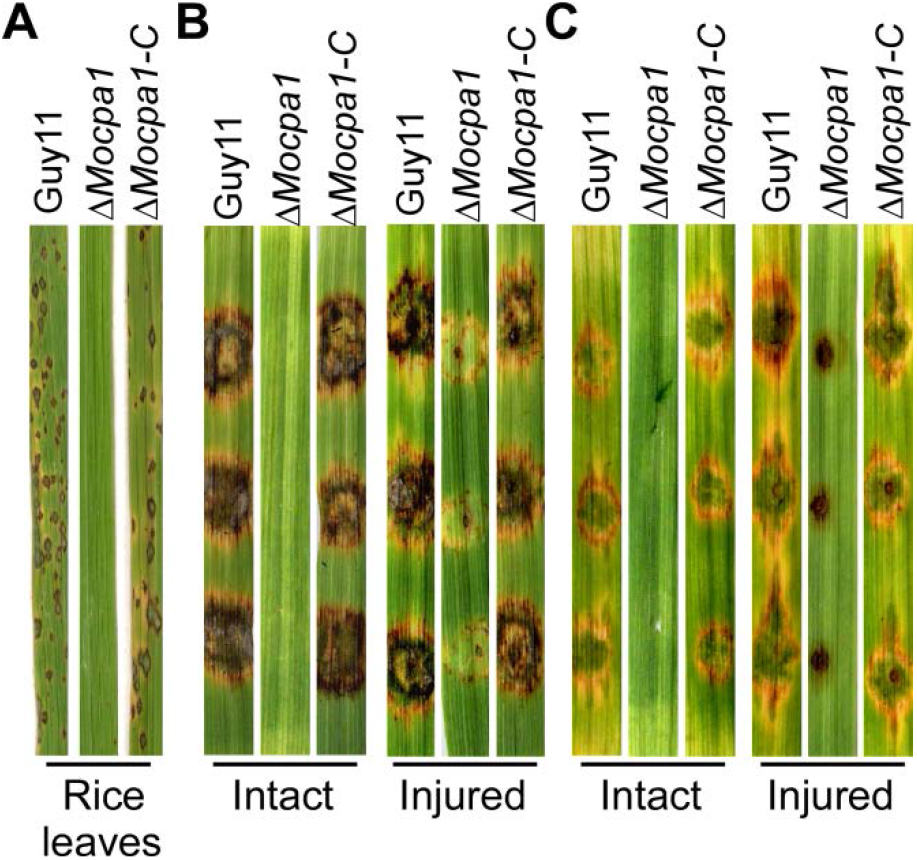
*MoCPA1* is critical for pathogenicity. (**A**) Spray inoculation assay of rice seedlings. Conidial suspensions (5×10^4^ spores/mL) were sprayed onto 3-week-old rice seedlings. Diseased leaves were imaged at 7 days after inoculation. (**B**) Mycelial plugs from WT and Δ*Mocpa1* mutants were inoculated onto intact and injured barley leaves imaged at 7 days. (**C**) Conidial suspensions (10 µL, 5×10^4^ spores/mL) were inoculated onto intact photographs were taken after 7 days of inoculation.

### Appressorium penetration and infectious hyphal growth are impaired in the Δ*Mocpa1* mutant

To determine the reason for the attenuated virulence of the Δ*Mocpa1* mutant, we first inoculated barley leaves and rice sheaths with conidia from the WT, Δ*Mocpa1* mutant, and the complemented strains and observed the resulting appressorium formation, appressorium penetration, and subsequent infectious hyphal growth. The results showed that, at 48 hours post-inoculation (hpi), the Δ*Mocpa1* appressoria were completely unable to penetrate the barley cells and the rice sheath and approximately 30% penetrated after 72 hpi. (Fig. 7A, 7B, and 7C). In addition to appressoria formed from conidia, *M. oryzae* is also known to develop appressorium-like structures by hyphal tips (Kim et al., 2009); thus, we examined the formation of the hypha-driven appressorium-like structures on barley leaves inoculated with mycelial plugs from the same set of strains. Unlike the WT and the complemented strain, which formed more appressorium-like structures, Δ*Mocpa1* formed very few appressoria from its hyphae (Fig. 7D). Further examination of the invasive hyphae revealed that, unlike the Guy11 and the complemented strain whose invasive hyphal growth occurred at 24 hpi, the Δ*Mocpa1* invasive hyphae developed at 72 hpi (Fig. 7C) and they could not colonize the neighboring cells (Fig. 7A and 7B). Based on these findings, we conclude that MoCpa1 plays a key role in conidium-mediated appressorium formation, hypha-driven appressorium formation, host penetration, and infectious hyphal growth and thus contributes immensely to *M. oryzae* pathogenicity.

**Fig. 7.**
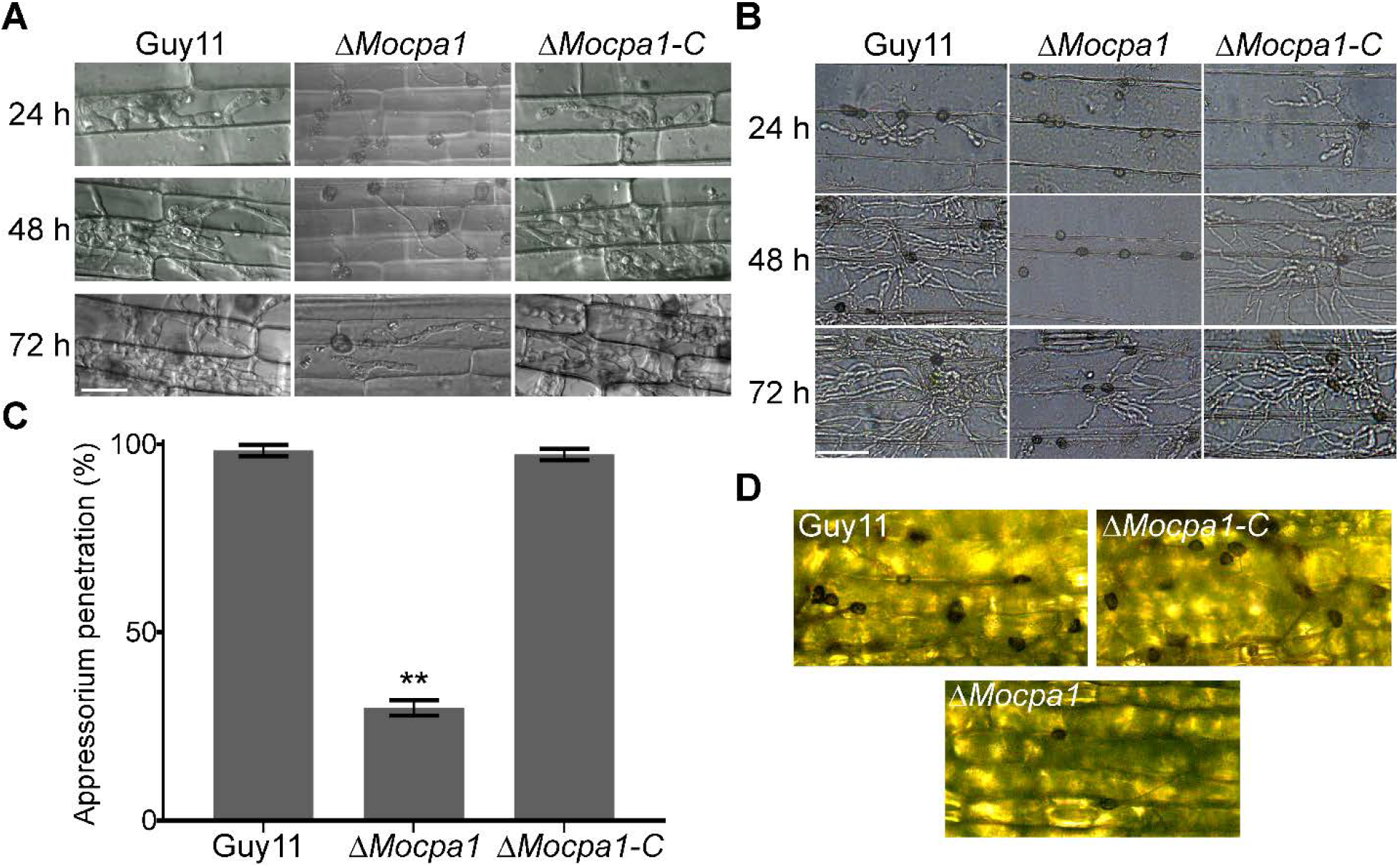
MoCpa1 is important for appressorium penetration and infectious hyphal development (**A, B**). The infection structures formed on rice and barley epidermal cells in response to inoculation with conidia of the wild-type Guy11, Δ*Mocpa1*, and complemented strains onto rice and barley leaves at 24, 48, and 72 h (A, rice leaves; B, barley leaves). Bar= 20 µm. **(C**) Statistical analysis of the percentage of appressorium penetration into barley leaves. Analysis of data was by use of GraphPad-prism6, error bars represent the standard deviation and the double asterisks (*) represent significant differences (p < 0.01) according to ordinary one – way ANOVA. (**D**). Barley leaf assay for hypha-driven appressorium formation. Mycelial plugs of Guy11, Δ*Mocpa1* mutant, and complemented strains were inoculated onto barley leaves, and hypha-mediated appressorium formation was observed at 24 h.

### Exogenous arginine partially rescues defects in conidiation and pathogenicity in Δ*Mocpa*

Since Δ*Mocpa1* exhibited arginine autotrophy, as shown in (Fig. 3C), we reasoned that the reduced sporulation and loss of pathogenicity in the Δ*Mocpa1* mutant may be attributed to arginine deficiency. To test this idea, we cultured the Δ*Mocpa1* mutant in sporulation rice bran media supplemented with 2.5 mM and 5 mM arginine. The spores obtained from the 2.5 mM arginine-supplemented cultures were then used to investigate whether pathogenicity defects observed with Δ*Mocpa1* mutant could be remediated on rice and barley leaves. Our results showed that exogenous arginine partially but not fully rescued the conidiation defects exhibited by Δ*Mocpa1* mutant (Fig. 8A and B). However, exogenous arginine failed to revert to the 3-three normal cell conidia and they were similar to those displayed in (Fig. 5A and 5B). Pathogenicity test with spores derived from 2.5 mM arginine-supplemented cultures also revealed a partial restoration of virulence by Δ*Mocpa1* mutant both on rice and barley leaves (Fig. 8C and 8D). These results demonstrate that conidiation and pathogenicity processes in *M. oryzae* require de novo arginine biosynthesis and that exogenous arginine cannot totally substitute for de novo arginine sources in *M. oryzae* to achieve maximum conidiation and pathogenicity efficiency.

**Fig. 8.**
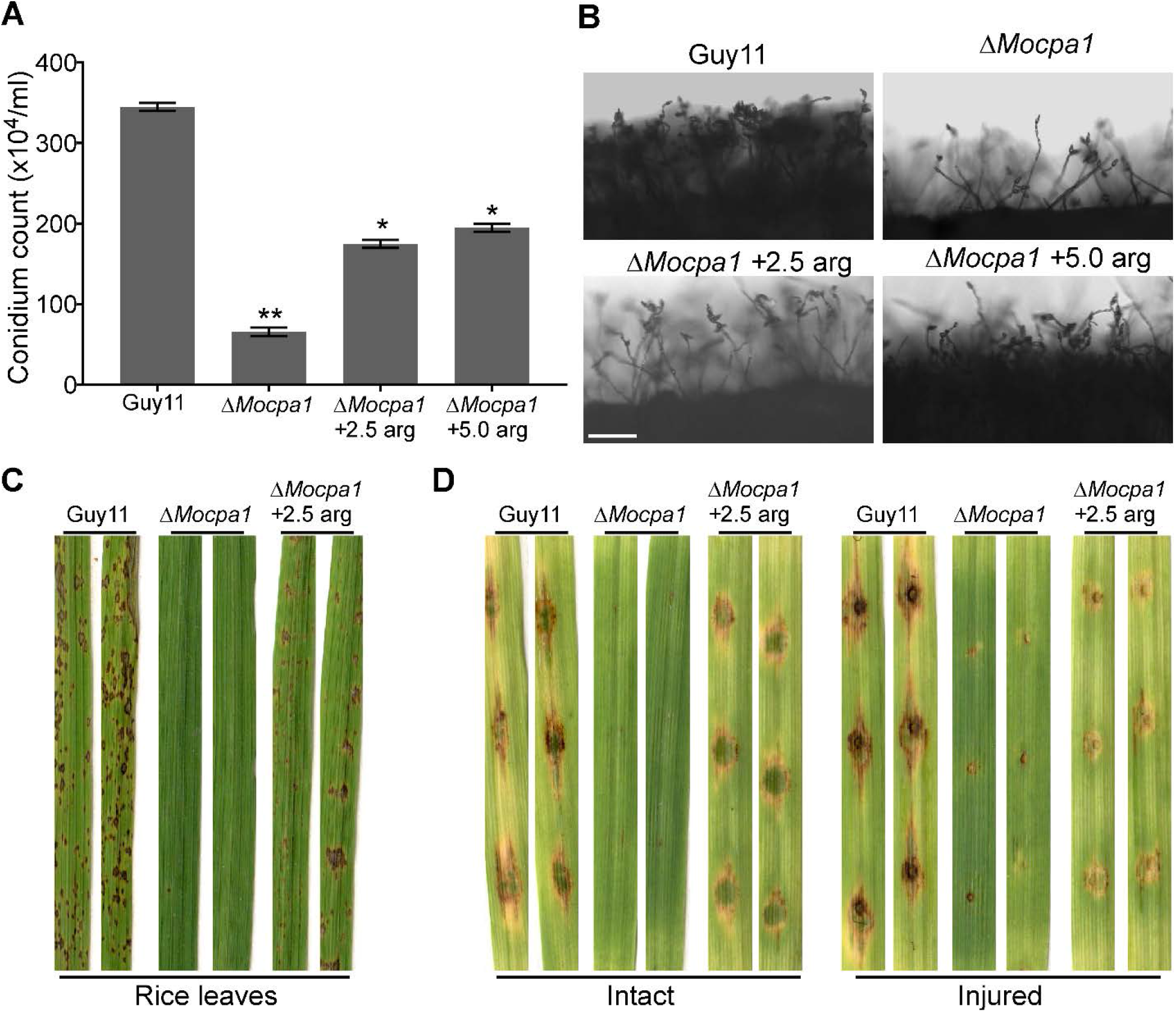
Exogenous arginine partially rescues defects in conidiation and pathogenicity of Δ*Mocpa1* strain. (**A**) Statistics showing the conidiation characteristics of Δ*Mocpa1* compared with the Guy11 wild-type strain. Compared with the WT strain, Δ*Mocpa1* was cultured in rice bran media supplemented with 2.5 mM arginine. Data analysis was by use of GraphPad-prism6 and Microsoft Excel spreads. Error bars represent the standard deviation and single and double asterisks (*) represent significant differences (p < 0.05 and p < 0.01) according to ordinary one – way ANOVA (**A**) Development of conidia on conidiophores observed on hydrophobic cover slips with a light microscope at 24 h after conidiation induction. Scale bar = 50 μm. (**C**and **D**) Pathogenicity of rice seedlings and barley leaves respectively. Conidia (1×10^5^ /mL) were harvested from 2.5 mM arginine-supplemented cultures, and additional 2.5 mM exogenous arginine was added into the conidial suspensions and then sprayed them onto 3-week-old rice seedlings, or dropped onto10-day-old barley leaves, and then photographed at 7 dpi.

### Deletion of *MoCPA1* caused reduced expression of rice pathogenesis-related genes

To evaluate the effects of *MoCPA1* on the induction of host immunity, we analyzed the expression of rice pathogenesis-related genes (*PRIa*, *PRIb*, *AOS2*, and *PBZ1*) using qRT-PCR upon rice leaf infection with conidia from the WT and Δ*Mocpa1* strain. Spores (5×10^4^ spores/mL) from the WT and Δ*Mocpa1* strain were sprayed onto 3-week-old rice seedlings and RNA extracted at 48 hpi. The Δ*Mocpa1* mutant elicited a relatively low host immunity compared to WT strain as evidenced by the reduced expression of both rice PR genes analyzed (Fig. 9).

**Fig. 9.**
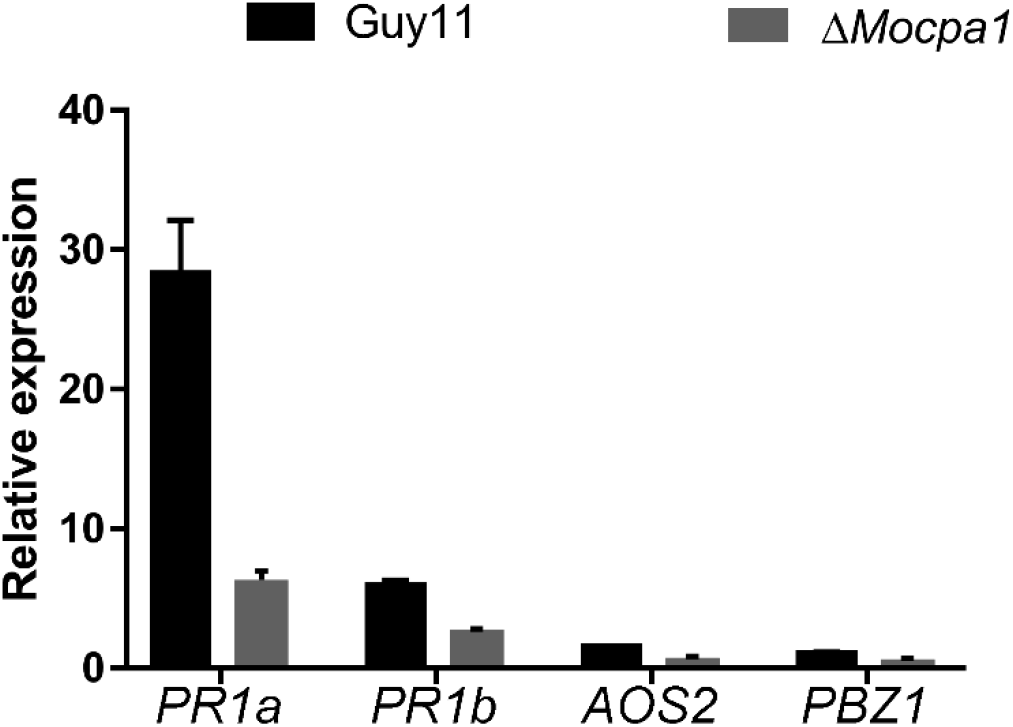
Compared with the Guy11 strain, the Δ*Mocpa1* failed to elicit a strong plant defense responses. Three-week-old rice leaf leaves were bombarded with spores of Guy11 and Δ*Mocpa1*, after which RNA was extracted at 48 hpi. The expression of the rice pathogenesis-related genes (*PR1a*, *PR1b*, *AOS2*, and *PBZ1*) was analyzed by qRT-PCR and normalized against the expression of the rice actin-encoding gene *OsACT2*. Δ*Mocpa1* induced a relatively low host immune response relative to the Guy11. Rice leaves inoculated with water (mock) was used as the control. Results are the average of three independent biological replications. The error bars represent the standard deviations.

### MoCpa1 is involved in the stress response

To investigate whether MoCpa1 contributes any role in fostering cell membrane and cell wall integrity, we cultured the Δ*Mocpa1* mutant, WT, and complemented strains in CM supplemented with a stress-reducing agent (DTT) and osmotic stress-inducing agent (NaCl) for 10 days and measured the colony diameter. There was no significant difference observed between the growth of the Δ*Mocpa1* mutant, Guy11, and the complemented strain on the CM plates supplemented with NaCl. However, there was impaired development of aerial hyphae for Δ*Mocpa1* in DTT (Fig. 10A). In addition, we also monitored the growth of the Guy11, Δ*Mocpa1*, and complemented strains on CM plates supplemented with cell wall integrity-enforcing agents including 200 μg/mL congo red (CR), 0.01% sodium dodecyl sulfate (SDS), and 200 μg/mL calcofluor white (CFW). Compared to the WT and the complemented strain, the Δ*Mocpa1* mutant was slightly inhibited in CR and SDS and highly inhibited in CFW (Fig. 10A 10B). These results demonstrate that MoCpa1 likely plays an essential role in stabilizing cell wall integrity in the rice blast fungus.

**Fig. 10.**
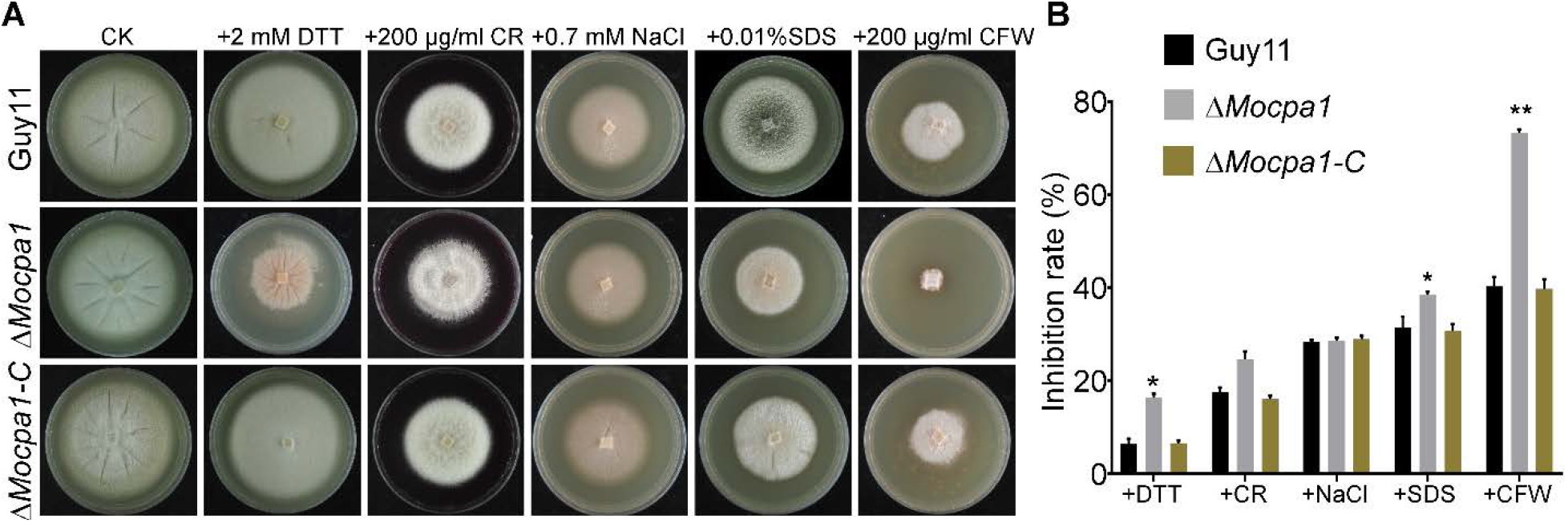
*MoCPA1* is involved in the stress response. (A) Colony morphology and diameter of the wild-type Guy11 strain and Δ*Mocpa1* mutant grown at 28°C under various stress conditions, 2 mM DTT, 0.7 M NaCl, 200 μg/mL Calcofluor White (CFW), 0.01% SDS or 200 μg/mL Conge Red (C.R.). Images were taken at 10 dpi. (B) Statistical representation of the response of Δ*Mocpa1* deletion mutants, the wild-type and the complemented strain to different cell membrane/wall stress inducing agents. The inhibition data was generated from three independent biological experiments with four technical replicates each time. One-way ANOVA (non-parametric) statistical analysis was carried out with GraphPad-prism6 and Microsoft Excel spreadsheet and error bars represent the standard deviation. Inhibition rate = (the diameter of untreated strain − the diameter of treated strain)/(the diameter of untreated strain) × 100%. Asterisks represent significant difference between the WT and the mutant (\**P*<0.05; \*\**P*<0.01)

### MoCpa1 localizes in mitochondria in *M. oryzae*

To determine the cellular compartmental localization of the MoCpa1 protein, a fluorescent reporter gene (eGFP) was fused to the C-terminus of the *MoCPA1* gene together with its native promoter and then cloned into a pKNTG vector containing a neomycin resistance gene. The construct was subsequently cotransformed into protoplasts of the WT strain with a pTE11 vector containing the mitochondrion-indicating marker (MoATP1). The MoCpa1-eGFP fusion proteins and mitochondrial ATP1-mCherry colocalized in the mitochondria in the growing hyphae, conidia, and appressoria (Fig. 11), thus indicating that MoCpa1 localizes to the mitochondria.

**Fig. 11.**
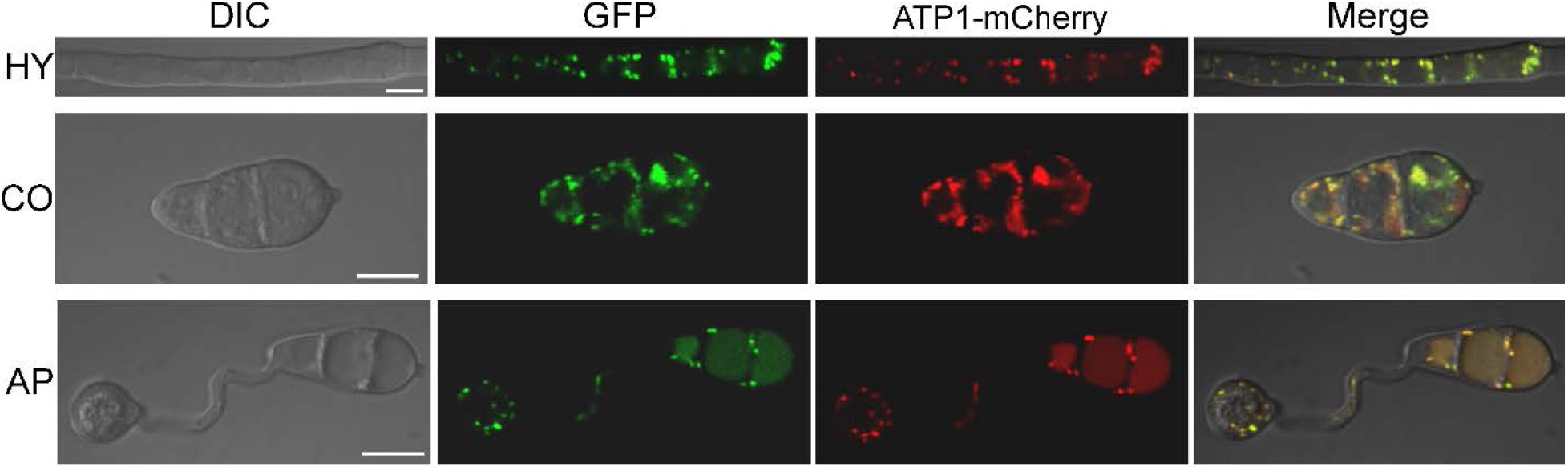
Subcellular localization of the carbamoyl phosphate synthase arginine-specific small subunit (MoCpa1) in *M. oryzae*. Hyphal (HY), Conidial (CO), and appressorium (AP) colocalization of MoCpa1-GFP together with the MoATP1-mCherry mitochondrial marker. For HY, bar = 20 μm. For CO and AP, bar = 50 μm.

## Discussion

In this study, we identified and functionally characterized the carbamoyl phosphate synthentase small subunit MoCpa1, in rice blast fungus. Bioinformatic analysis revealed that MoCpa1 is a 471-amino acid protein containing a CPSase_sm_chain domain and a GATase domain. Further Cpa1 domain prediction from other fungi confirmed CPA1 is highly conserved among the several filamentous fungi and might have conserved functions in fungal development or pathogenicity. Deletion of *MoCPA1* resulted in impairment of aerial hyphae formation, conidiation, appressorium formation and penetration, hyphal appressorium-mediated formation, stress response, hyphal appressorium-formation, and pathogenicity. Previous study showed that amino acid biosynthesis is important for the fungal infection cycle (Xin Liu et al., 2019; Orasch et al., 2019; Tan, Zhao, He, Zhang, & Yi, 2020). In *M. oryzae*, arginine biosynthesis catalyzed by MoArg1, MoArg5, MoArg6, MoArg7, and MoCpa2 have been shown to be indispensable for the development and pathogenicity of *M. oryzae* (Xinyu Liu, Cai, et al., 2016; Y. Zhang et al., 2015). Our growth assay showed attenuated growth of Δ*Mocpa1* deletion mutants on MM medium, with the growth defects being rescued upon addition of exogenous arginine, clearly demonstrating the importance of MoCpa1 in arginine biosynthesis. This results were in agreement with a recently reported study on the role of *CPA1* in glutamine biosynthesis in fungal pathogen *Colletotrichum gloeosporioides* (Tan et al., 2020).

Nutrient deficiency had been shown to be one of the contributors to autolysis in fungi (White, McIntyre, Berry, & McNeil, 2002). Increased expression and repression of genes involved in autolysis in autolytic and nonautolytic cells has been reported in *Aspergillus nidulans* (Emri et al., 2006). In this study, the Δ*Mocpa1* mutant showed poor development of aerial hyphae on OTM and SDC media. Analysis of expression of autolysis-related genes, MGG_07877, MGG_13610, MGG_13185, MGG_00086, MGG_01247, MGG_05533, and MGG_01001 by qRT-PCR detected upregulation of these genes in the Δ*Mocpa1* deletion mutant, demonstrating a likelihood of Δ*Mocpa1* mutant cells undergoing autolytic degradation caused by arginine starvation.

Conidia play an important role in the disease cycle of rice blast (Wilson & Talbot, 2009). Investigation of asexual reproduction revealed that the Δ*Mocpa1* strain was compromised in terms of sporulation, presenting a 75% reduction in conidium production compared to that of WT and the complemented strain. Although exogenous arginine partially rescued the conidiation defects in the Δ*Mocpa1* deletion mutant, the conidia number were still half the number of the WT, demonstrating that de novo arginine biosynthesis mediated by *MoCPA1* is crucial for conidiogenesis in the rice blast fungus. Evaluation of the expression of conidiation-related genes revealed that the expression of genes was reduced in the Δ*Mocpa1* mutant, implying that the reduced conidiation in Δ*Mocpa1* could be attributed to the downregulation of these genes. Our findings were consistent with those of a previous study showing that reduced sporulation of the Δ*Moppe1* mutant was linked to the downregulation of conidiation-related genes (Qian et al., 2018). These results, therefore, indicate that *MoCPA1* is important for conidiation in *M. oryzae*. Further examination of Δ*Mocpa1* conidial morphology revealed that, in addition to the normal three-cell conidia, the Δ*Mocpa1* deletion mutants produced 10% and 40% conidia comprising one cell and two cells, respectively, thus confirming further that *MoCPA1* is also crucial for conidial morphogenesis.

In rice blast fungus, it was reported that deletion of carbamoyl phosphate synthase arginine specific large subunit *MoCpa2*, leads to attenuated virulence (Xinyu Liu, Cai, et al., 2016). Moreover, a recently reported study, showed that loss of *CPA1* in *Colletotrichum gloeosporioides* resulted in complete loss of virulence (Tan et al., 2020). Similarly, the Δ*Mocpa1* mutant was nonpathogenic on barley and rice leaves. The explanation for the attenuated virulence exhibited by Δ*Mocpa1* mutant was due to defects in hypha-driven appressorium formation, conidium-mediated appressorium formation, appressorium penetration, and invasive hyphal growth, all of which are critical processes for infection of *M. oryzae*. Introduction *MoCPA1* gene in Δ*Mocpa1* deletion mutant completely restored these phenotype while addition of exogenous arginine partially rescued the pathogenicity defects of Δ*Mocpa1*, thus demonstrating the importance of MoCpa1-mediated arginine biosynthesis in infection-related morphogenesis in *M. oryzae*.

Expression of pathogenesis-related (PR) genes upon fungal infection is one of the indicators for studying the host defense response. The host defense response to rice blast fungus through the expression of PR genes has previously been reported (Guo et al., 2010; Xinyu Liu, Qian, et al., 2016). In our study, analysis of the rice PR genes (*PR1A*, *PR1B*, *PPZ1*, and *AOS1*) upon rice infection by the WT and Δ*Mocpa1* mutant spores resulted in the Δ*Mocpa1* mutant eliciting a weaker host immune response as evident from lower PR gene expression compared to the WT strain. This results were consistent with those of previous study in which the SAICAR synthetase null mutant Δ*Moade1* was nonpathogenic and failed to elicit a strong plant defense response (Fernandez, Yang, Cornwell, Wright, & Wilson, 2013). The relatively low induction of rice immunity by the Δ*Mocpa1* mutant shows that the host immune response was not the main reason for the defects in invasive hyphal growth and colonization; rather, arginine deficiency was the main reason, as evident from the partial restoration of pathogenicity of the Δ*Mocpa1* deletion mutant for both barley and rice leaves upon the addition of exogenous arginine.

Localization assays revealed that MoCpa1 is mitochondrial bound, supporting its role in the synthesis of carbamoyl phosphate, a reaction that has been reported to occur in mitochondria (Shi, Caldovic, & Tuchman, 2018).

In summary, we have demonstrated that MoCpa1 is involved in arginine biosynthesis and is essential for aerial hyphal growth, conidiation, conidial morphogenesis, appressorium formation, host penetration, invasive hyphal growth and pathogenicity in rice blast fungus. The findings provide further evidence on the importance of amino acid biosynthesis in the development of pathogenic fungi. Since Cpa1 are conserved in fungi, these proteins may act as a suitable target for exploitation as a novel antifungal target

## Material and methods

### Fungal strains and culture conditions

*M. oryzae* Guy11 was used as the wild-type (WT) strain and was used to generate Δ*Mocpa1* deletion mutants. To promote their vegetative growth, the WT strain and mutant were cultured at 25°C using complete media (CM: 0.6% yeast extract, 0.6% casein hydrolysate, 1% sucrose, 1.5% agar) as previously described (J. Chen et al., 2008). Other media used in this study included minimal media (MM: 6 g of NaNO_3_, 0.52 g of KCl, 0.52 g of MgSO_4_, 1.52 g of KH_2_PO_4_, 10 g of glucose and 15 g of agar in 1 L of double-distilled water), straw decoction and corn media (SDC: 100 g of rice straw, 40 g of corn flour, and 15 g of agar in 1 L of double-distilled water) and oatmeal agar media (OMA: 50 g of oatmeal and 15 g of agar in 1 L of double-distilled water).

For fungal protoplast preparation and transformation, previously described standard protocols were adopted (H.-L. Wang, Kim, Siu, & Breuil, 1999; Wendland, 2003). Screening of hygromycin- and neomycin-resistant transformants was performed on TB3 media supplemented with 250 μg/mL hygromycin B (Roche Applied Science) and 200 μg/mL G418 (Invitrogen), respectively. Genomic DNA and total RNA were extracted from mycelia cultured in liquid CM with shaking at 110 rpm at 28°C for 3 days. For conidiation assays, the strains were cultured on rice bran agar media for 10 days at 28°C (2% rice bran, 1.5% agar; pH 6.5) at 28°C for 10 days in the dark followed by 3 days of continuous light illumination.

The conidia were collected in 5 m of distilled water, filtered through three layers of lens paper and counted by using a hemocytometer under a microscope. Conidium germination and appressorium formation on the hydrophobic surfaces were measured as described previously (Qi et al., 2012).

### Target gene deletion and complementation in *M. oryzae*

To generate Δ*Mocpa1* mutants, a split-marker homologous recombination approach was used (Tang et al., 2015). The entire *MoCPA1* gene was replaced with the hygromycin gene (*HPH*). The upstream and downstream flanking sequences of the *MoCPA1* gene were amplified from *M. oryzae* genomic DNA and used to generate split markers using splicing overlap extension by fusing with upstream and downstream hygromycin sequences amplified from a pCX62 vector. Primers used in this study are found at (Table S1).

For complementation, a 1.6-kb full-length open reading frame (ORF) together with its 2.4-kb native promoter was amplified using primers Cpa1com F/ Cpa1com R (Table S1) and cloned behind the green fluorescent protein (GFP) in a pKNTG vector containing a neomycin resistance gene to generate a pKNTG-MoCPA1-GFP fusion cassette, which was later transformed into Δ*Mocpa1* mutant protoplasts. The complemented strains were screened by neomycin resistance and observations of GFP fluorescence.

### Appressorium formation, penetration and infection assays

Conidia harvested from 10-day-old rice bran agar cultures, were filtered through three layers of lens paper, and were resuspended to a final concentration of (5×10^4^ spores/mL) using double distilled water. For host infection assays spores, were adjusted to 1.5–2.0 × 10^5^ conidia/mL in 0.02%v/v Tween-20 solution and sprayed onto 21-days-old rice seedlings. Plants were incubated in a humid chamber at 28°C for 24 h in darkness. They were then transferred to another humid chamber that had a 12-h photoperiod. Disease severity was assessed by taking images of 6-cm-long rice blades 7 days after inoculation. For the barley infection assay, 10 μL of conidial suspension (5×10^4^ spores/mL) was added onto the barley leaves, after which the plants were incubated at 28°C for 24 h in darkness. They were then transferred to lighted conditions and imaged after 7 days

For appressorium formation assays, 20 μL of conidial suspension was added to an artificial hydrophobic coverslip and incubated in darkness at 26°C. Appressorium formation was then examined at 4 h, 8 h, and 12 h and 24 h time intervals Barley penetration assays were performed by inoculating 10 μL droplets of conidial suspension (5×10^4^ spores/mL) repeatedly onto 10-day-old barley seedlings. Penetration and invasive hyphal development were examined at 24 h, 48 h, and 72 h.

To examine appressorium-like structure and mycelial-based penetration, culture blocks of 4-day-old strains, grown on solid CM were inoculated onto 10-day-old barley leaves for 24 hrs and penetration and appressorium-like structure observed according to the previously described method (Kong et al., 2013).

For the rice sheath assay, 100 μL of conidial suspension (5×10^4^ spores/mL) was inoculated into the inner leaf sheath cuticle cells. The plants were then incubated for 24 hrs 48 hrs and 72 hrs at 28°C under humid conditions and the leaf sheaths were observed under a microscope as described previously (L. Zhang et al., 2011).

### Nucleic Acid Manipulation, Southern Blotting Analysis, and qRT-PCR

To extract genomic DNA, strains were cultured in liquid CM medium for 3-4 days, mycelia were filtered then frozen in liquid nitrogen, all the strains genomic DNA were extracted according to the Cetyltrimethylammonium bromide (CTAB) method described by (Y. Wang et al., 2017)

For Southern blot assays, an open reading frame (ORF) probes for the *MoCPA1* gene and the hygromycin (HPH) gene were amplified with primer pairs CPa1 koF/ CPa1 koR and Hph F/ Hph respectively (Table S1). Probe labeling, hybridization, and detection were performed with a DIG High Prime DNA Labeling and Detection Starter Kit (Roche Applied Science, Penzberg, Germany). Total RNA was isolated from frozen fungal mycelia using an RNA extraction kit (Megan, China).

To measure the relative abundance of *MoCPA1* transcripts during various stages of fungal development, the method described (Dong et al., 2015) was adopted. Total RNA samples were extracted from mycelia grown in liquid CM, from conidia, and from plants inoculated with Guy11 conidia (1 × 10^8^ spores/mL) for 8, 24, 48, and 72 h.

For the evaluation of the transcript level of PR genes, RNA samples were extracted from the rice leaves inoculated with Guy11, with the Δ*Mocpa1* mutant (5×10^4^ spores/mL), and with water (mock) for 48 h.

To measure the transcription of conidiation and autolysis related genes, RNA was extracted from Guy11 and the Δ*Mocpa1* mutant cultured in liquid CM and shaken for 3 days at 110 rpm. All the RNA extracted in this study was reverse transcribed using SYBR^®^ Premix Ex Taq™ (Tli RNaseH Plus) (Takara Biomedical Technology, Beijing Co., Ltd, China). An Eppendorf Realplex2 Mastercycler (Eppendorf AG 223341, Hamburg, Germany) was used to perform qRT-PCR.

### Microscopy

An Olympus DP80 light microscope (Japan) was used to observe conidiophore development, conidium shapes, appressoria on hydrophobic surfaces, appressorium penetration, and invasive hyphal development. For the confocal microscopy assay, a confocal microscope equipped with a Nikon A1 plus instrument (Nikon, Tokyo, Japan) was used to observe the fluorescence of Guy11-expressing MoCpa1-GFP and mitochondrial ATP1-mCherry.

### Construction of domain architecture and phylogenetic analysis

BLASTp using *S. cerevisiae* Cpa1 as query was used to search for the MoCpa1 in the *M. oryzae* genome database http://www.kegg.jp/kegg-bin/show_organism?org=mgr. Amino acid sequences of all the other fungi were acquired from the National Center for Biotechnology Information (NCBI), https://www.ncbi.nlm.nih.gov for the domain prediction. Prediction of the subcellular localization was performed by the use of PSORT II (http://psort.hgc.jp/), and domains were predicted in Pfam (http://pfam.janelia.org/). Cpa1 domain sequence were represented by construction of domain architecture using Ibis1.0 program (W. Liu et al., 2015). Alignment and phylogenetic analysis of the obtained amino acid sequences were performed using MEGA version 6, while phylogeny was generated using the maximum-likelihood method, with branches of the tree tested with 1000 bootstrap replicates.

### Statistical analysis

At least three replicates of each result was presented as the mean standard deviation (SD). The significant differences between the WT and the mutant tests were statistically evaluated by SD and one-way analysis of variance (ANOVA) using Graph prism6. The data between specific different tests were compared statistically by ANOVA, followed by F-test if the ANOVA result is significant at P* < 0.05, P **< 0.01

## Supplementary materials

Supplementary material 1 *MoCPA1* gene deletion strategy and Southern blot confirmation

**Figure S1.**
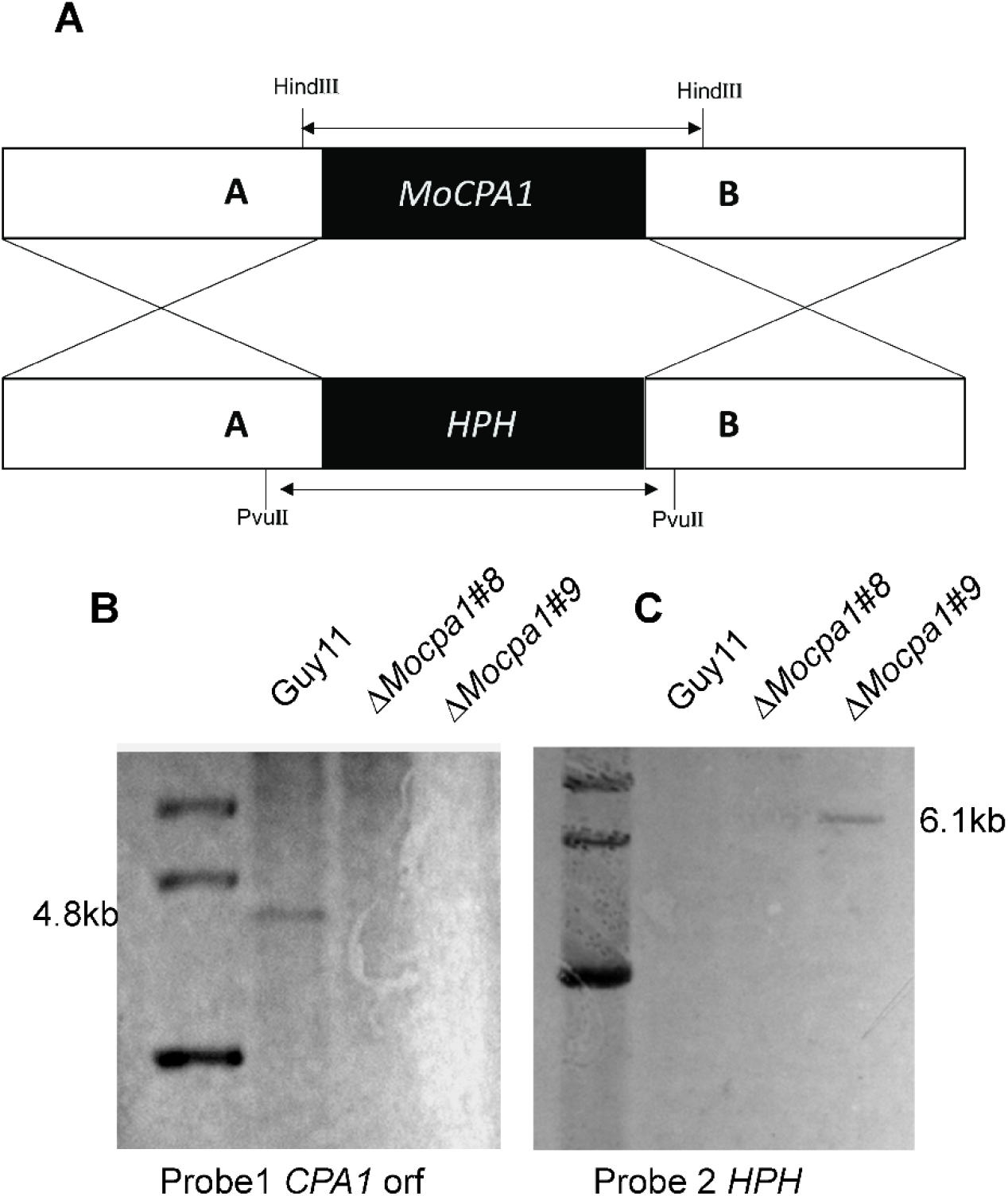
Southern blot analysis of the *MoCPA1* gene deletion mutant. (**A**) Strategy of knocking out MoCPA1 in the *M. oryzae* genome. (**B**) Southern blot analysis of the gene knockout mutants and WT Guy11 via *MoCPA1* ORF probe 1 (**C**) Southern blot analysis of the gene knockout mutants and WT Guy11 via hygromycin phosphotransferase (hph) (probe hph).

**Figure S2.**
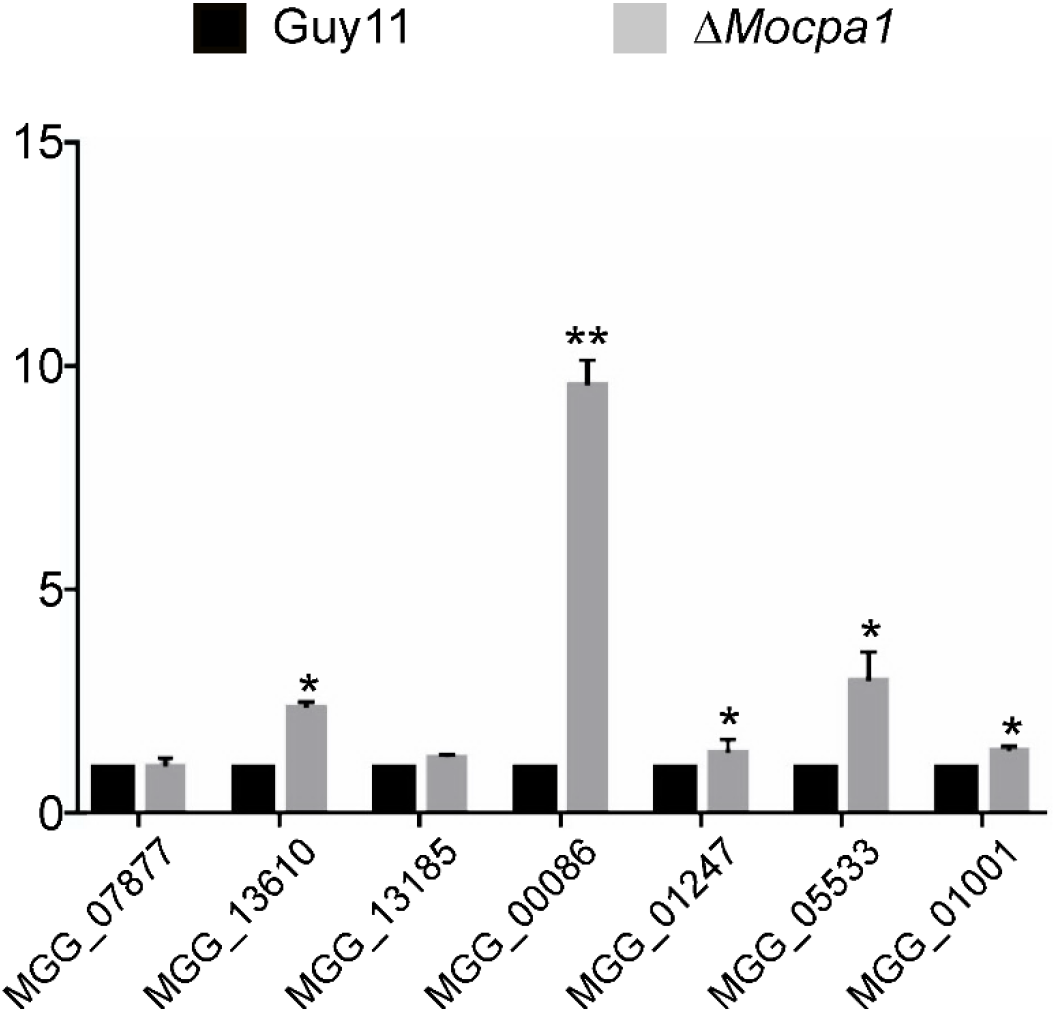
Deletion of *MoCPA1* resulted in upregulataion genes involved in autolysis. RNA was extracted from WT and Δ*Mocpa1* mutant cultured in liquid CM with shaking for 3 days at 110 rpm. The *ACTIN* gene (MGG_03982) was used for normalization. Data was analyzed with GraphPad-prism6, the error bars represents the standard deviation and the asterisks (\**P*<0.05) and (\*\**P*<0.01) represent significance deference in expression based on one way-ANOVA

**Table S1.**
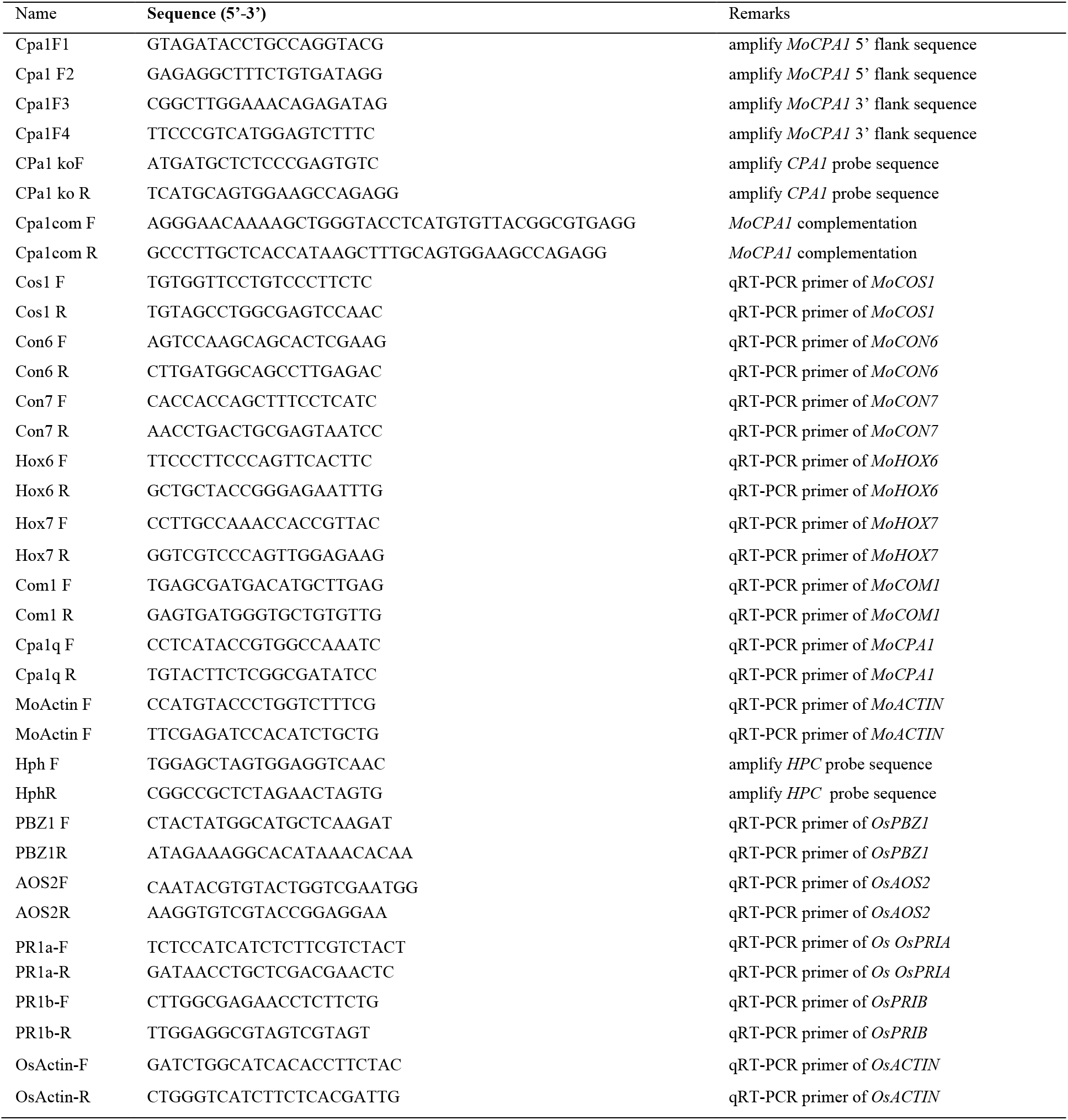
Oligonucleotide primers used in this study.

## Acknowledgment

This research was supported by the National Key R and D program of China (2016YD0300700) to Z. W. We are grateful to Justice Norvienyeku and Professor Stefan Olsson for advice and discussion on the project

## Notes

### Competing Interest Statement

The authors have declared no competing interest.

